# Lipopolysaccharide (LPS) induces increased epidermal green autofluorescence of mouse

**DOI:** 10.1101/501189

**Authors:** Yujia Li, Mingchao Zhang, Yue Tao, Weihai Ying

**Affiliations:** Med-X Research Institute and School of Biomedical Engineering, Shanghai Jiao Tong University, Shanghai 200030, P.R. China; Collaborative Innovation Center for Genetics and Development, Shanghai 200043, P.R. China

**Keywords:** Inflammation, LPS, Autofluorescence, Skin, Keratin 1

## Abstract

Our recent studies have suggested that characteristic ‘Pattern of Autofluorescence (AF)’ of each disease could be a novel biomarker for non-invasive diagnosis of multiple major diseases such as acute ischemic stroke. It is necessary to determine if increased epidermal green AF may be produced by major pathological factors such as inflammation. In our current study, we used C57BL/6Slac mice exposed to LPS to test our hypothesis that inflammation may induce increased epidermal green AF: LPS rapidly induced significant increases in the epidermal green AF of the mice’s ears at 1 hr after LPS injection. LPS also dose-dependently increased the epidermal green AF. The AF intensity had a linear relationship with the LPS dosages at both 3 and 7 days after the LPS administration. The AF images exhibited the characteristic structure of the keratinocytes in *Stratum Spinosum*, suggesting that the origin of the increased AF was keratin 1 and/or keratin 10. Collectively, our current study has provided the first evidence indicating that inflammation can rapidly and dose-dependently induce increased epidermal green AF, suggesting that the green AF may be the first biomarker for non-invasive and rapid detection of systemic inflammation. Since inflammation is a key pathological factor of numerous diseases, our finding has highlighted the value of the epidermal AF as a novel diagnostic biomarker for numerous diseases.

## Introduction

A number of studies have indicated that autofluorescence (AF) of skin or blood can be used for non-invasive diagnosis of diabetes (14) and cancer (22). Our recent study has indicated that UV-induced epidermal green AF could be a novel biomarker for predicting UV-induced skin damage (8). The epidermal green AF appears to be originated from UV-induced keratin 1 degradation (8), which is mediated by UV-induced oxidative stress (10).

Our recent studies have suggested that characteristic ‘Pattern of AF’ could be a novel biomarker for non-invasive diagnosis of acute ischemic stroke (2), acute myocardial infarction (24), stable coronary artery disease (24), Parkinson’s disease (3) and lung cancer (11). While oxidative stress could be a key factor leading to the increased epidermal green AF of the patients’ skin, future studies are necessary to determine if the increased AF may also be caused by other pathological factors such as inflammation.

Inflammation is a crucial pathological factor in multiple major diseases including cerebral ischemia (18), acute myocardial infarction (16), stable coronary artery disease (9), Parkinson’s disease (15) and lung cancer (12). Because inflammation can lead to increased levels of cytokines and oxidative stress (7), we proposed our hypothesis that inflammation may also induce increased epidermal green AF. In our current study, we used mice exposed to lipopolysaccharide (LPS) to test this hypothesis. Our study has suggested that inflammation could be an important factor that produces the increased epidermal green AF, which may be used as the first biomarker for non-invasive and rapid determinations of systemic inflammation.

## Materials and Methods

All chemicals were purchased from Sigma (St. Louis, MO, USA) except where noted.

### Animal Studies

Male C57BL/6Slac mice, ICR mice, and BALB/cASlac-nu nude mice of SPF grade were purchased from SLRC Laboratory (Shanghai, China). All of the animal protocols were approved by the Animal Study Committee of the School of Biomedical Engineering, Shanghai Jiao Tong University.

### Administration of LPS

Male C57BL/6Slac mice with the weight of 20 – 25 g were administered with 0.1, 0.5, or 1 mg/kg LPS with intraperitoneal (i.p.) injection. The stock solution of LPS with the final concentration of 0.2 mg/ml was made by dissolving LPS in PBS. The mice were administered with these doses of LPS every 24 hrs.

### Imaging of epidermal AF

As described previously (8), a two-photon fluorescence microscope (A1 plus, Nikon Instech Co., Ltd., Tokyo, Japan) was used to image the epidermal AF of the ears of the mice, with the excitation wavelength of 488 nm and the emission wavelength of 500 – 530 nm. The AF was quantified as follows: Sixteen spots with the size of approximately 100 × 100 μm^2^ on the scanned images were selected randomly. After the average AF intensities of each layer were calculated, the sum of the average AF intensities of all layers of each spot was calculated, which is defined as the AF intensity of each spot.

### Western blot assays

Western blot assays were conducted as described previously (8): The lysates of the skin were centrifuged at 12,000 *g* for 20 min at 4°C. The protein concentrations of the samples were quantified using BCA Protein Assay Kit (Pierce Biotechonology, Rockford, IL, USA). As described previously(20), a total of 50 μg of total protein was electrophoresed through a 10% SDS-polyacrylamide gel, which were then electrotransferred to 0.45 μm nitrocellulose membranes (Millipore, CA, USA). The blots were incubated with a monoclonal Anti-Cytokeratin 1 (ab185628, Abcam, Cambridge, UK) (1:4000 dilution) or actin (1:1000, sc-58673, Santa Cruz Biotechnology, Inc., Dallas, TX, USA) with 0.05% BSA overnight at 4°C, then incubated with HRP conjugated Goat Anti-Rabbit IgG (H + L) (1:4000, Jackson ImmunoResearch, PA, USA) or HRP conjugated Goat Anti-mouse IgG (1:2000, HA1006, HuaBio, Zhejiang Province, China). An ECL detection system (Thermo Scientific, Pierce, IL, USA) was used to detect the protein signals. The intensities of the bands were quantified by densitometry using Image J.

### Statistical analyses

All data are presented as mean ± SEM. Data were assessed by one-way ANOVA, followed by Student – Newman-Keuls *post hoc* test. *P* values less than 0.05 were considered statistically significant.

## Results

One hr, 1 day, 3 day and 7 day after the mice were administered with 1 mg/kg LPS by i.p injection, the skin’s green AF of the mice’s ears was determined. LPS induced significant increases in the green AF at all of the time points (Figs. 1A and 1B). We also determined the effects of 0.1, 0.5 and 1 mg/kg LPS on the green AF, showing that LPS dose-dependently induced increases in the green AF, assessed at 3 (Figs. 2A and 2B) and 7 days (Figs. 3A and 3B) after the LPS administration. The AF intensity has linear relationship with the LPS dosages at both 3 days (R^2^ = 0.9697) and 7 days (R^2^ = 0.9889) after the LPS administration. These AF images exhibited the characteristic structure of the keratinocytes in *Stratum Spinosum*.

**Figure 1.**
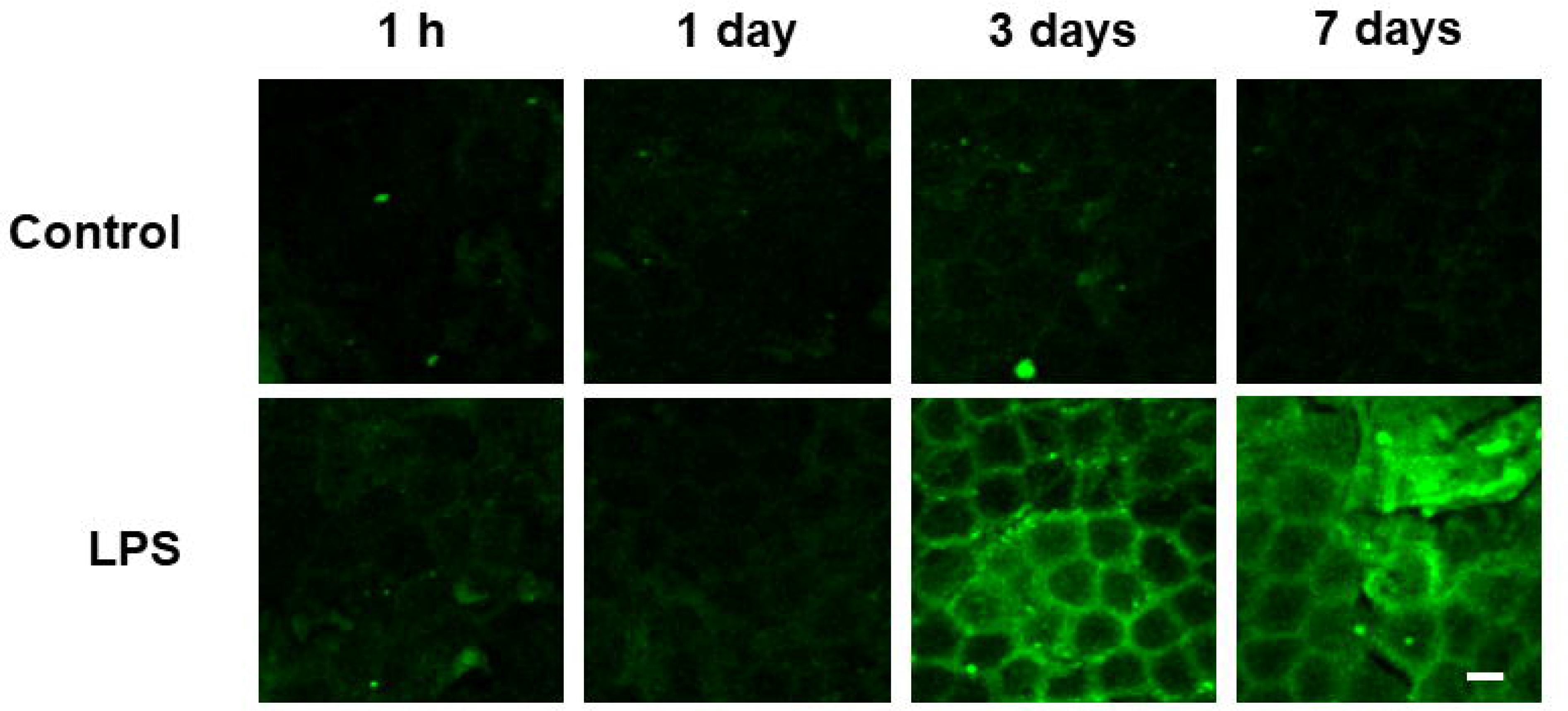
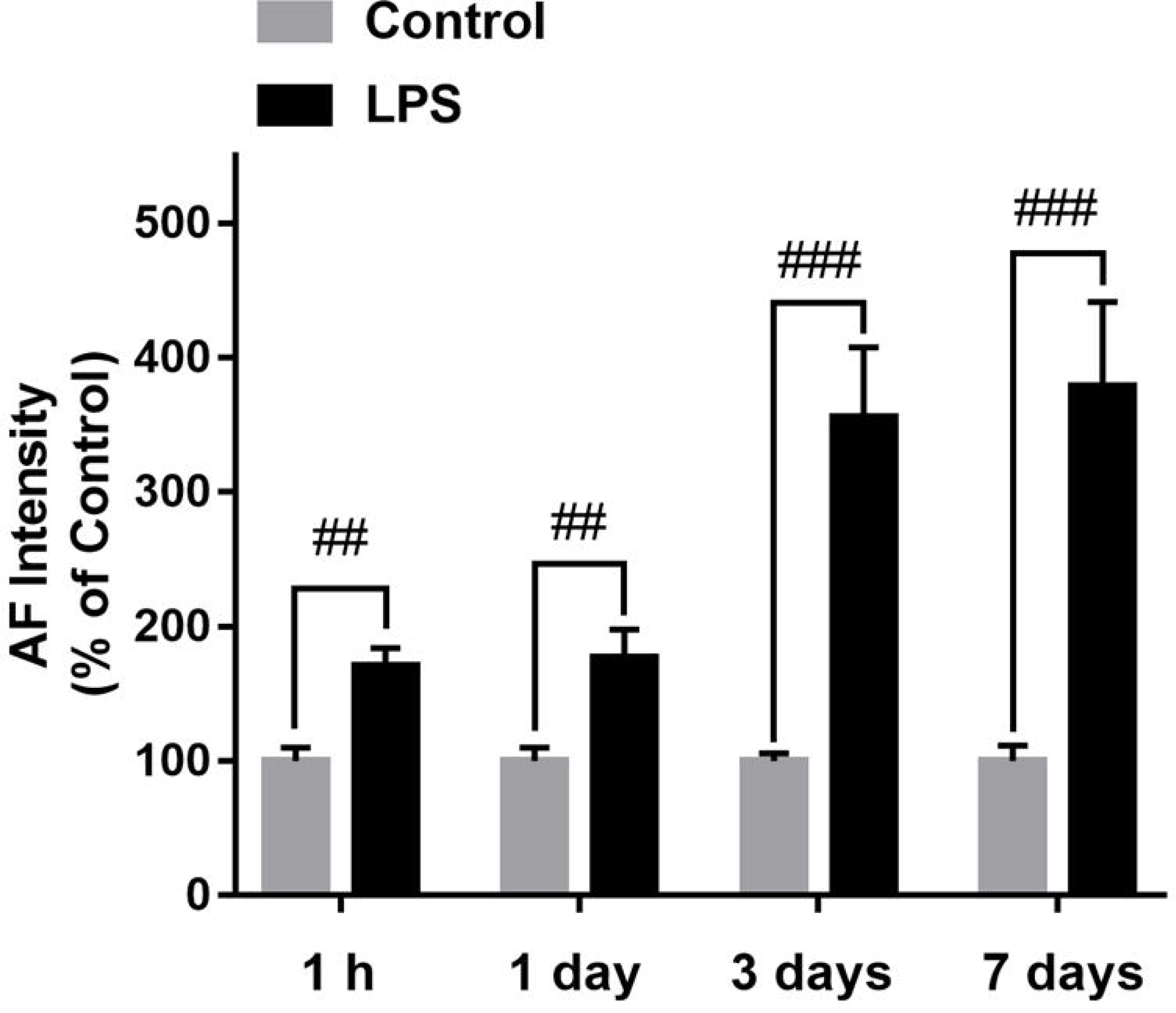
LPS rapidly induced significant increases in the epidermal green AF of C57 mouse’s ears. (A) Intraperitoneal (i.p) injection of 1 mg/kg LPS increased the skin’s green AF of the ears, assessed at 1 hr and 1, 3 and 7 days after the LPS injection. Excitation wavelength = 488 nm and emission wavelength = 500-530 nm. Scale bar = 20 μm. (B) Quantifications of the AF showed that 1 mg/kg LPS induced significant increases the skin’s green AF of the ears, assessed at 1 hr and 1, 3 and 7 days after the LPS injection. N = 6 – 15. **, *P* < 0.01; ***, *P* < 0.001.

**Figure 2.**
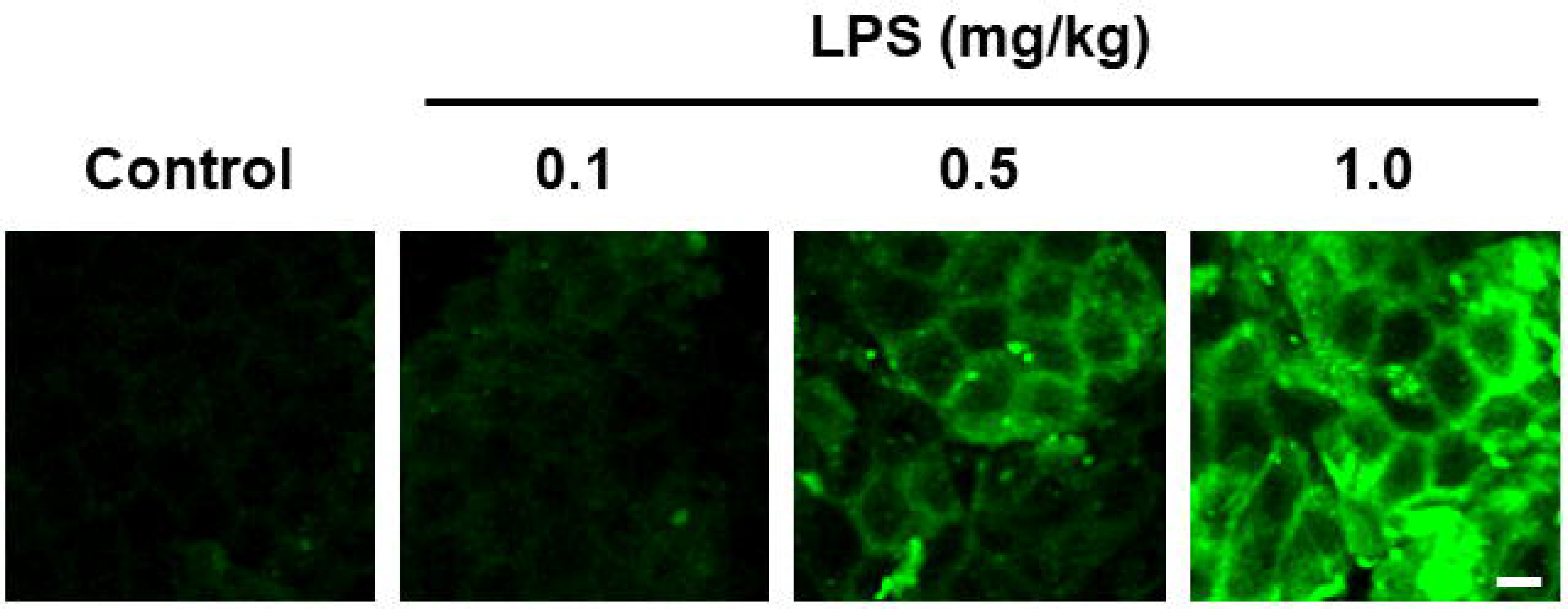
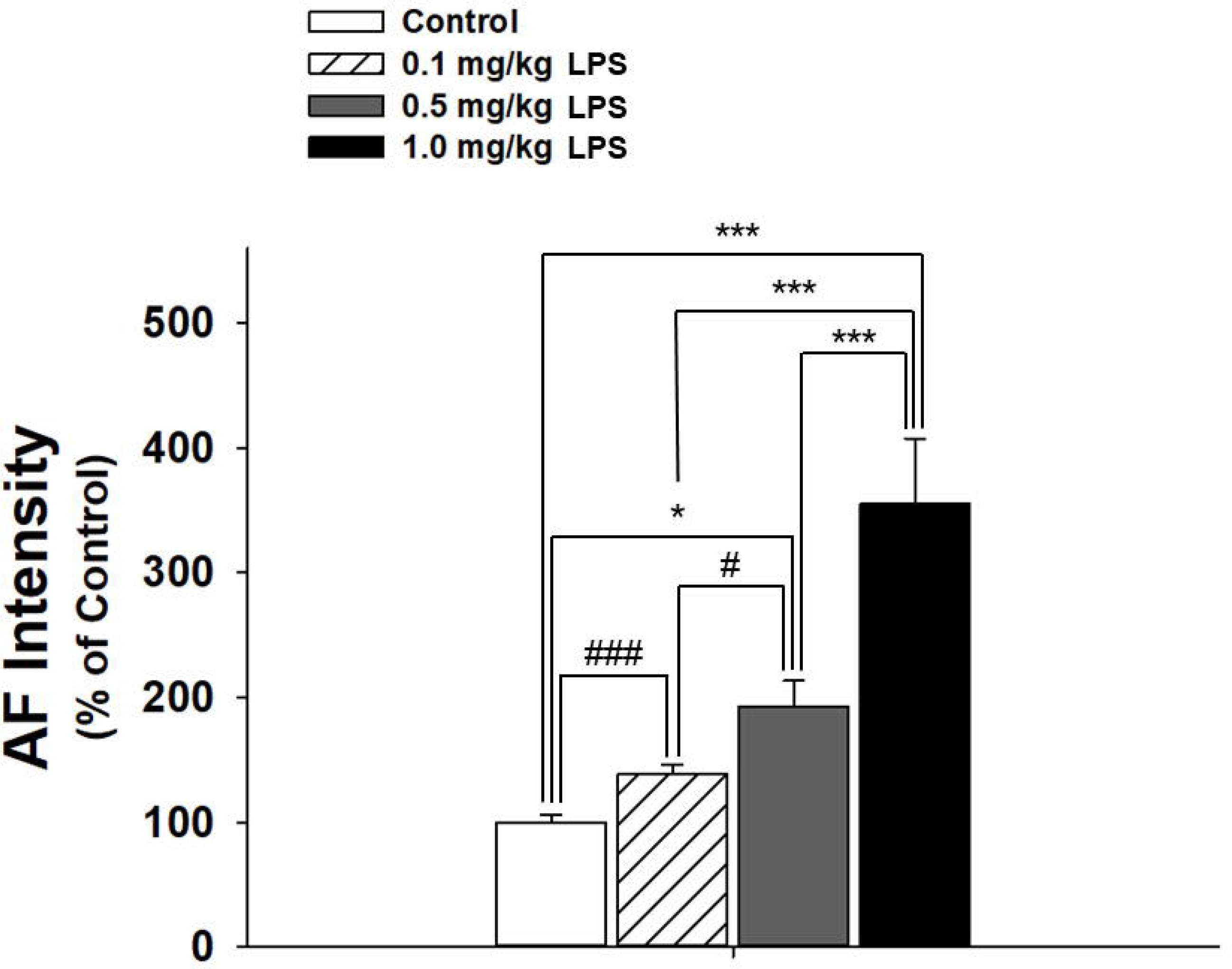
LPS dose-dependently induced increases in the epidermal green AF of C57 mouse’s ears 3 days after LPS injection. (A) Intraperitoneal injection of 0.1, 0.5 and 1 mg/kg LPS does-dependently increased the skin AF of the ears, assessed at 3 days after the LPS injection. Excitation wavelength = 488 nm and emission wavelength = 500-530 nm. Scale bar = 20 μm. (B) Quantifications of the AF showed that 0.5 and 1 mg/kg LPS induced significant increases in the skin AF of the ears. Three days after mice were administered with 0.1, 0.5 and 1 mg/kg LPS, the epidermal green AF images of the mice’ ears were taken under a two-photon fluorescence microscope. N = 6. ##, *P* < 0.01; ###, *P* < 0.001 (Student *t*-test).

**Figure 3.**
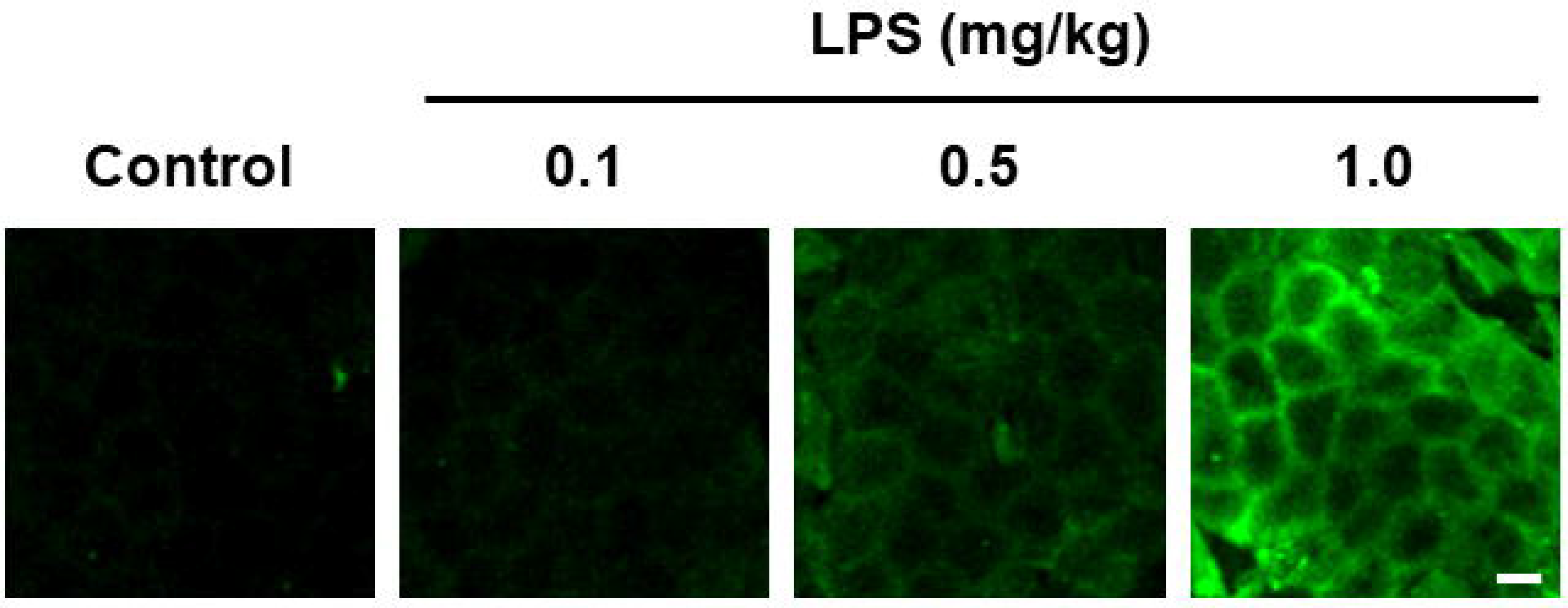
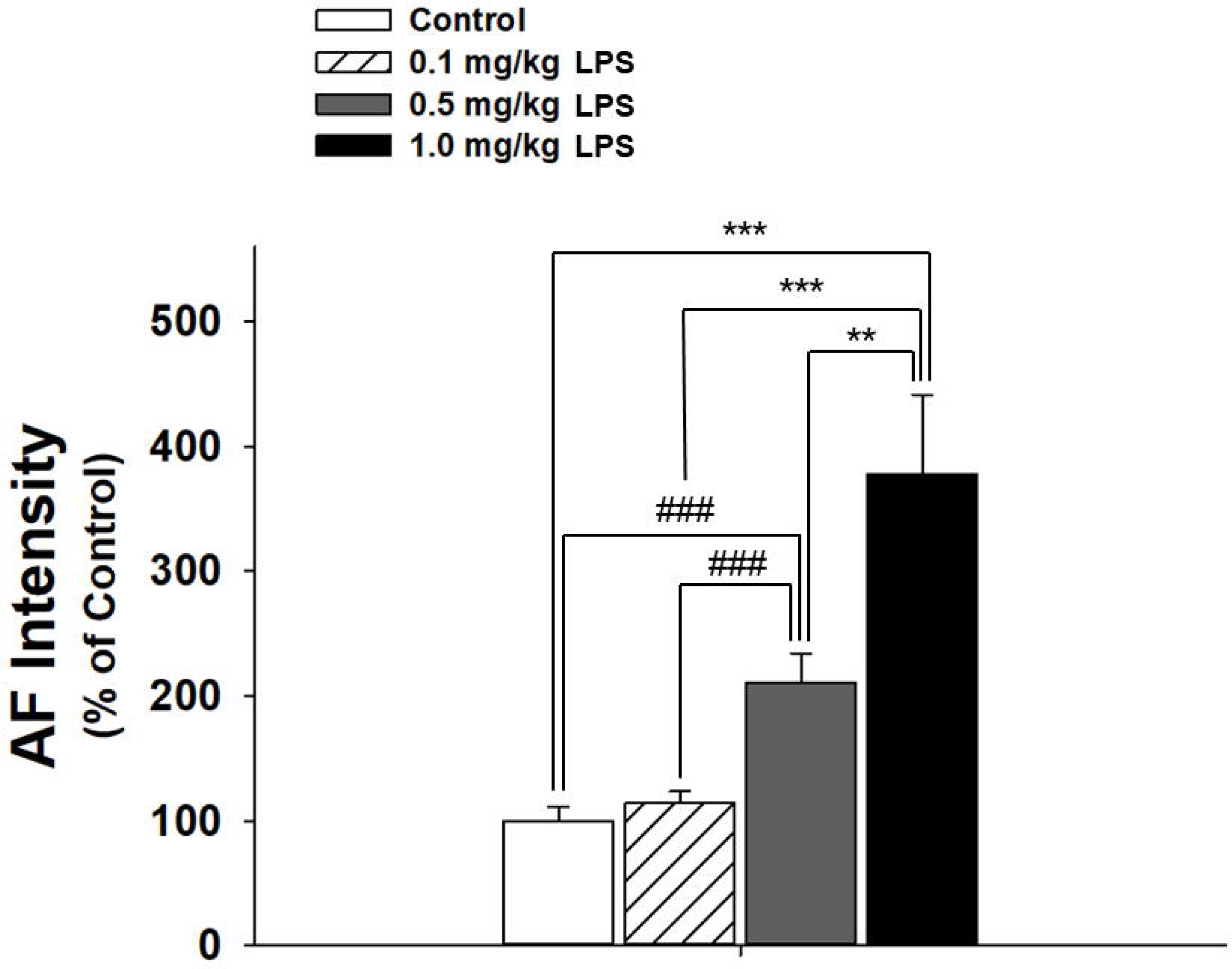
LPS dose-dependently induced increases in the epidermal green AF of C57 mouse’s ears 7 days after LPS injection. (A) Intraperitoneal injection of 0.1, 0.5 and 1 mg/kg LPS does-dependently increased the skin AF of the ears, assessed at 7 days after the LPS injection. Excitation wavelength = 488 nm and emission wavelength = 500-530 nm. Scale bar = 20 μm. (B) Quantifications of the AF showed that 0.5 and 1 mg/kg LPS induced significant increases in the skin AF of the ears. Seven days after mice were administered with 0.1, 0.5 and 1 mg/kg LPS, the epidermal green AF images of the mice’ ears were taken under a two-photon fluorescence microscope. N = 6. ##, *P* < 0.01; ***, *P* < 0.001.

Seven days after the mice were administered with 0.1, 0.5 and 1 mg/kg LPS, the keratin 1’s (Krt 1’s) level of the ears’ skin was determined by Western blot. We found that LPS led to a significant decrease in the keratin 1 level of the ears’ skin (Figs. 4A and 4B).

**Figure 4.**
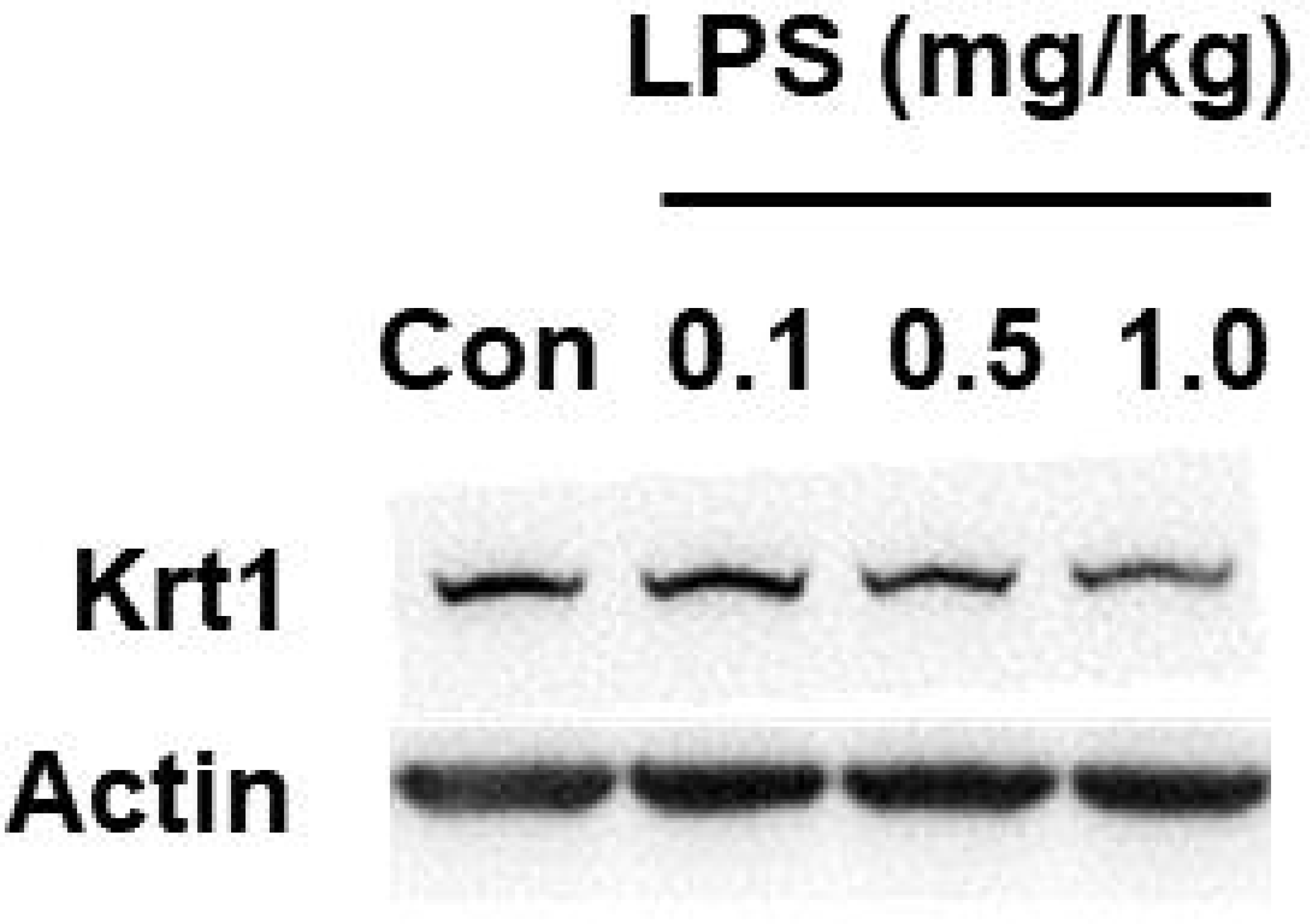
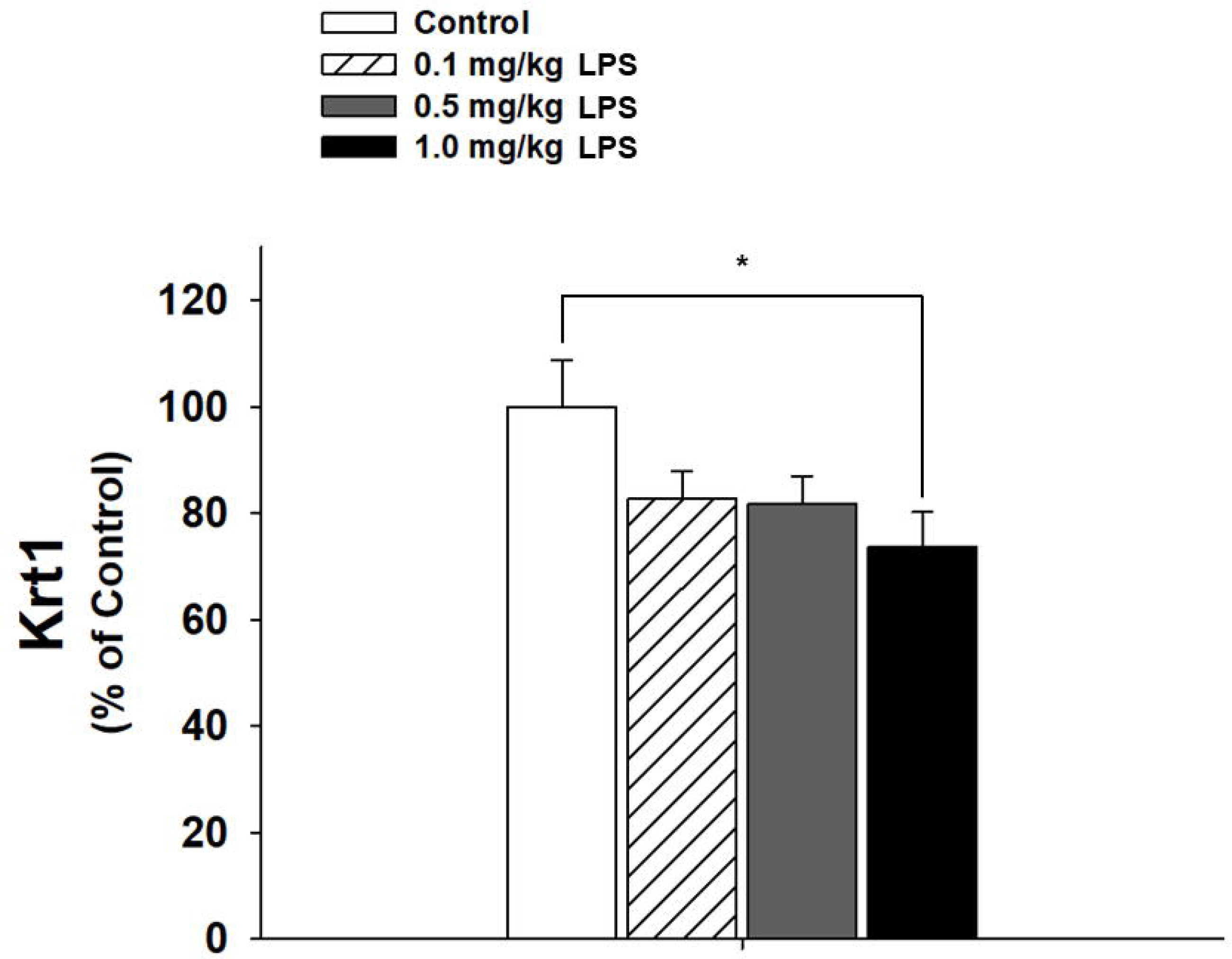
LPS administration led to Krt 1 degradation of C57 mouse’s ears. (A) Western blot assays showed that i.p injection of 1 mg/kg LPS led to a decrease in the Krt 1 level of the ears, assessed at 7 days after the LPS administration. (B) Quantifications of the Western blot showed that 1 mg/kg LPS led to a significant decrease in the Krt 1 level of the ears. Seven days after mice were administered with 0.1, 0.5 and 1 mg/kg LPS, the Krt 1’s level of the ears were determined by Western blot. *, *P* < 0.05. N = 12.

## Discussion

The major findings of our study include: First, LPS rapidly induced significant increases in the epidermal green AF of mice’s ears at 1 hr after LPS injection; second, the AF intensity was highly correlated with the LPS dosages; and third, the AF images exhibited the characteristic structure of the keratinocytes in *Stratum Spinosum*.

As shown by our study, inflammation can rapidly and dose-dependently induce increased epidermal green AF, suggesting that the green AF may be used as a novel biomarker for rapid detection of systemic inflammation. Inflammation is a key pathological factor of numerous diseases (9,12,16,18). However, so far there has been no non-invasive method for determining the levels of systemic inflammation. Our findings have suggested that the epidermal green AF may become the first biomarker for non-invasive determinations of systemic inflammation as well as numerous inflammation-associated diseases.

Our recent studies have suggested that characteristic ‘Pattern of AF’ could be a novel biomarker for non-invasive diagnosis of acute ischemic stroke (2), myocardial infarction (24), stable coronary artery disease (24), Parkinson’s disease (3) and lung cancer (11). While oxidative stress could be a key factor leading to the increased epidermal green AF of the patients’ skin (10), our current study has suggested that inflammation could be another important factor that produces the increased AF intensity. The characteristic state of the inflammation in these major diseases may produce the characteristic increases in the epidermal green AF in the patients’ body, thus forming the characteristic ‘Pattern of AF’.

Our study has found that the structure of the LPS-induced green AF, similar with the UV-induced green AF (8), exhibited the characteristic polyhedral structure of the keratinocytes in *Stratum Spinosum*. This finding has suggested that the LPS-induced green AF was located in the keratinocytes in *Stratum Spinosum*. Because the only epidermal fluorophores that selectively exist in the keratinocytes of *Stratum Spinosum* are Krt 1 and Krt 10 (4,13,21), our study has suggested that Krt 1 and/or Krt 10 are the origin of the LPS-induced increases in the epidermal green AF.

Keratins play multiple significant roles in epithelium, including intermediate filament formation (19), inflammatory responses (5,17) and cellular signaling (1,6). Krt 1 and its heterodimer partner Krt 10 are the major keratins in the suprabasal keratinocytes of epidermis (4,13,21), which is a hallmarker for keratinocyte differentiation (23). As found in UV-exposed mouse skin (10), our current study has also found that LPS induced Krt 1 degradation of the mouse’s ears 7 days after the LPS injection. Future studies are warranted to further identify the molecules responsible the LPS-induced epidermal green AF.

## Acknowledgments

The authors would like to acknowledge the financial support by a Major Research Grant from the Scientific Committee of Shanghai Municipality #16JC1400500 and #16JC1400502 (to W.Y.), and a Major Special Program Grant of Shanghai Municipality (Grant # 2017SHZDZX01) (to W.Y.),

